# Copy when uncertain: Lower light levels result in higher trail pheromone deposition and stronger reliance on pheromone trails in the ant *Lasius niger*

**DOI:** 10.1101/473579

**Authors:** Sam Jones, Tomer J. Czaczkes, Alan J. Gallager, Jonathan P. Bacon

**Affiliations:** School of Life Sciences, University of Sussex, Falmer BN19QG, UK; Department of Zoology and Evolutionary Biology, University of Regensburg, Regensburg 93053, Germany; Brighton and Sussex Medical School, University of Sussex, Falmer BN19QG, UK

**Keywords:** Light levels, information conflict, copy when uncertain, route learning, pheromone deposition, ants

## Abstract

Animals may gather information from multiple sources, and these information sources may conflict. Theory predicts that, all else being equal, reliance on a particular information source will depend on its information content relative to other sources. Information conflicts are a good area in which to test such predictions. Social insects, such as ants, make extensive use of both private information (e.g. visual route memories) and social information (e.g. pheromone trails) when attempting to locate a food source. Importantly, eusocial insects collaborate on food retrieval, so both information use and information provision may be expected to vary with the information content of alternative information sources. Many ants, such as *Lasius niger*, are active both day and night. Variation in light levels represents an ecologically important change in the information content of visually-acquired route information. Here, we examine information use and information provision under high light levels (3200 lux), moderate light levels simulating dusk (10 lux) and darkness (0.007 lux). Ants fail to learn the location of a food source in darkness. As light levels decrease, ants show decreasing reliance on private visual information, and increasing pheromone trail following, consistent with a ‘copy when uncertain’ strategy. In moderate light levels and darkness, pheromone deposition increases, presumably to compensate for the low information content of visual information. Varying light levels for cathemeral animals provides a powerful and ecologically meaningful method for examining information use and provision under varying levels of information content.

## Introduction

Strategic information use is critical to the success of many animals. Animals must, for example, decide whether to explore new options, exploit the knowledge they already have, or use information gleaned from or sent by other animals about potential options [1–3]. Two important classes of information for animals are private information and social information. Private information can only be accessed by the focal animal, and includes genetic information, internal states, and importantly, memories. Social information is information gathered from the behaviour of other animals [4]. These may be cues, such as evidence of conspecifics having recently been in a particular location [5,6], or intentionally produced social signals, such as the waggle dance of honey bees or pheromone trails deposited by ants [7,8].

While much research effort has been focussed on assessing whether to exploit available information or gather (often costly) new information [2,9], once information is gathered animals must decide how to use multiple information sources. Matters are complicated when conflicts arise between information sources. When this occurs, one option is to produce and follow a weighted intermediate value [10–12]. However, sometimes an intermediate response is not possible, for example when deciding between two feeding locations. Alternatively, a hierarchy of information sources can be employed, with one type of information being exclusively used until it is not available, after which others begin to be employed [13]. A more nuanced strategy is to weigh up the usefulness or information richness of different information sources, and follow the best one [14].

Social insects, such as ants and bees, offer an unique system in which to study information use strategies [1,3], as social information use is likely to be very well developed in this group. More fundamentally, however, in many aspects of information use by social insects no conflict is expected within a colony. For example, we expect no conflicts over which resource to use. This should lead to full honesty in communication, making social information more valuable. Critically, it also means that information providers, rather than being exploited, are benefiting from providing information. This in turn is expected to result not only in strategic information use in the receiver, but also in strategic information provision by the signaller. The context in which social insects choose to actively produce social information can be as informative as the context in which they choose to respond to it [1]. For example, ants which are more likely to make a mistake (and thus presumably are more uncertain) have been found to deposit less pheromone [15]. Conversely, ants which made a mistake when outgoing, but do eventually find the food source, deposit more pheromone than ants which initially make a correct decision [15,16]. Ants from colonies with poor learning performance also tend to deposit more pheromone when returning from a food source [17].

These considerations should, in principle, strengthen the effect of social information on the behaviour of social insects. It is therefore surprising that in most cases in which conflict between social signals and private information have been studied, ants and bees predominantly follow their own memories [8,18–27]. While this is not a universal pattern [19,28,29], it is nonetheless striking.

Reversals in information use, from reliance on memories to reliance on social signals, are informative for understanding this phenomenon. One reason for the neglect of social information is that social information is often information-poor when compared to memories. For example, the information about food quality conveyed by waggle dances, pheromone trails, or the presence of conspecifics is extremely noisy. By contrast, private memories of a food sources’ quality are very accurate. Insects may be attempting to follow a ‘copy-if-better’ strategy [2], but without accurate quality information, rarely copy. And indeed, when unambiguous quality information about a better food source is provided, *Lasius niger* ants switch from following memories to following pheromone trails [30]. Another important reversal in social information use was reported by Beugnon & Fourcassie [21,31]. During daylight hours and in a well-lit laboratory, *Formica pratensis* wood ants followed memories over pheromone trails. However, at night their behaviour reversed, and they were found to preferentially follow pheromone trails. This suggests that ants may also be following a ‘copy-when-uncertain’ strategy, only relying on chemical signals when memories are unavailable or unreliable. Such a strategy has been previously reported in other social insects in other contexts, such as flower choice in bumblebees [32] and during nest relocation in rock ants [33]. Reports also describe ants depositing pheromone to lower-quality resources only in the dark [34], or increasing pheromone deposition when learning was unsuccessful [15] or on hard-to-learn routes [16].

The use and provision of information by animals under different light regimes offers a promising means of studying information conflict and information use strategies. Variation in light levels over many orders of magnitude is a challenge many animals have to cope with every day-night cycle. This variation in light levels results in strong variation in the certainty of visual route memories – the main source of navigational information for many ants [35–37]. Here, we study the use and provision of social information under different light levels in the ant *Lasius niger*. Under high light levels, *L. niger* preferentially follow private route memories over pheromone trails, even if the pheromone trails are very strong [22,30]. We first test confirm that route memories in *L. niger* are based solely on visual cues (as reported by [36]). We then ask whether *L. niger* foragers modulate their pheromone deposition (social information production) in response to different light levels, and find that pheromone deposition increases as light levels decrease. Finally, we assess their preference for private information (memories) over social information (pheromone trails) at different light levels, and find a stronger reliance on social information as light levels decrease.

## Methods

### Study Species

Colonies of the black garden ant, *Lasius niger*, were collected from Falmer in East Sussex, UK. Each colony was housed in a plastic container (30 × 30 ×10 cm high) with a plaster of Paris base containing a circular nest cavity constructed from plaster of Paris (13.5 cm diameter × 1.5 cm high) and covered by a disc of dark card. All colonies were queenless with 1,000 – 3,000 workers and small numbers of brood. Queenless colonies readily forage, produce trails and are commonly used in behavioural experiments [36,38], remaining viable for 18 months or more. The ants were fed three times a week on a Bhatkar mix [39], with ad libitum access to water. To ensure motivation, feeding was stopped 4 days prior to experimentation.

### General experimental design

Following the method of Grüter et al. [22] we constructed a foraging trail as shown in Figure 1. A white cardboard bridge (20 × 2 cm) connected the colony container to a transparent polycarbonate plastic T-maze covered with white paper. The stem of the T was 15 cm long and each branch was 11 cm long, with a consistent width of 2cm. Experiment 1 was run in a small windowless room with an ambient temperature of 22 ^°^C. Experiments 2 and 3 were carried out in a small room containing various items of lab equipment and furniture which served as visual landmarks for the foraging ants. We used 3 light levels in our experiments: bright light (3100 lux), moderate light (10 lux), and darkness (0.0007 lux). Light intensity was measured repeatedly throughout the experiments using a photometer (LI-COR inc; model LI-188B) to ensure illumination was consistent within treatment replicates. These luminances were chosen to reflect normal daylight, crepuscular light, and a moonless night, respectively. A portable halogen work light (IP 44; model NXS-500P) with a 500 w halogen bulb was used to provide high intensity illumination for the bright light treatment and a floor lamp with a 230 w linear halogen bulb and dimmer switch (Dar; model OPU 4946) provided illumination for the moderate light treatment. In the moderate light and darkness experiments infrared light was used to provide illumination for experimental working and behavioural observations, but this long wavelength illumination was not detectable by the ants; as in humans, most insects have trichromatic vision [UV, blue and green in the case of insects; [40], but see [41] for evidence of bichromatic vision in an ant]. However, their visible spectrum is shifted towards shorter wavelengths than ours [41,42]; for example the spectral sensitivity maxima (λ_max_) for the ants *Atta sexdens* and *Camponotus blandor* are 500 nm and 570 nm respectively [41,43], considerably shorter than the 700 nm found in humans [44]. To provide infrared light a sleeve created from 2 ply corrugated cardboard tightly fitted over the hood of an angle poise lamp with a 60 w bulb. Two 50 mm square 665 nm long pass (IR) filters (Schott; model FRG-66550) were slotted tightly together into a hole cut in the centre of the cardboard hood so that when switched on the lamp only provided infra red light

**Fig 1.**
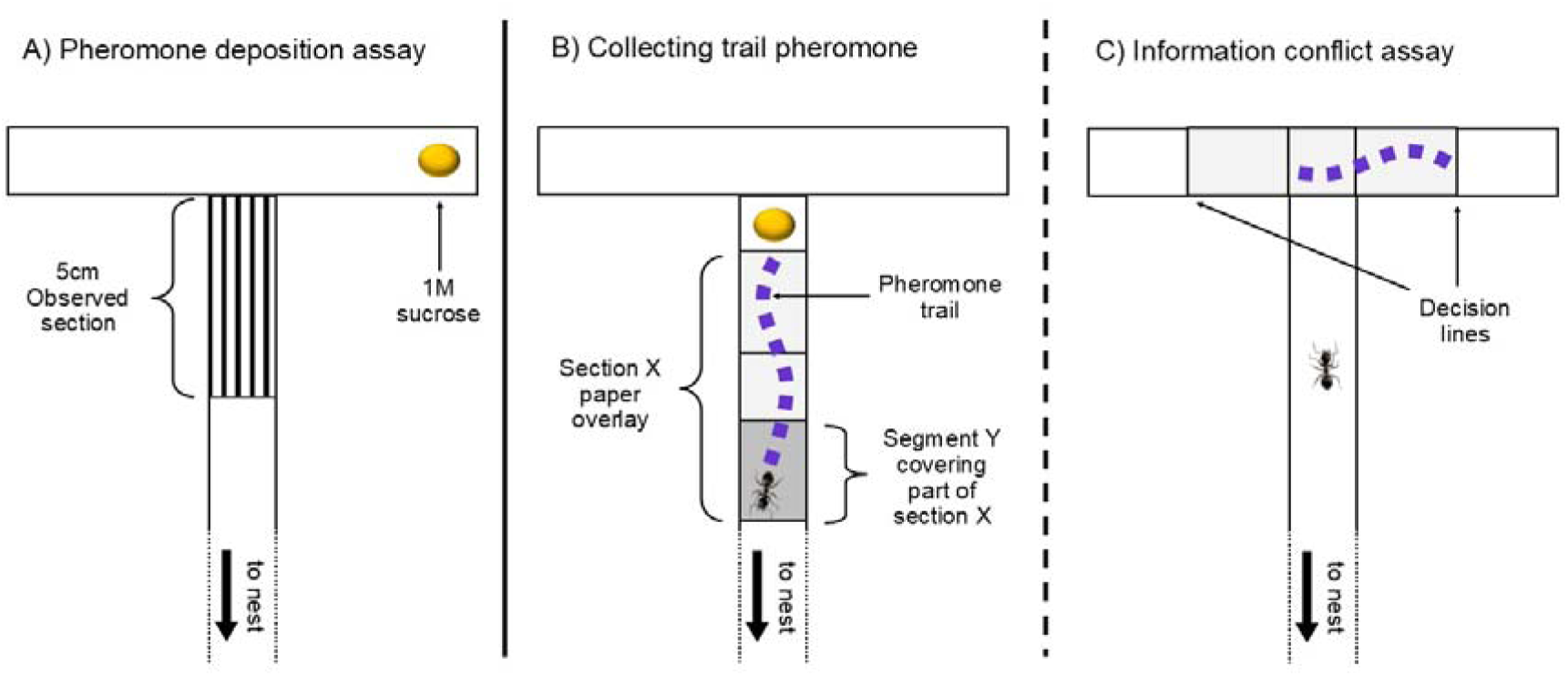
**A)** Experimental design used to measure trail pheromone deposition frequency by foraging ants under three different lighting regimes (darkness, moderate and bright). For each treatment frequency of deposition was recorded for the observed section (lined area) for three journeys: the first return journey to the nest, the first outward to the food source and the second return to the nest. Each experiment involved 8-12 ants that were marked with paint while feeding on the sugar solution for the first time. **B)** Experimental design used to acquire trail pheromone for subsequent conflict situations. A section of paper (X) was partially covered by a segment (Y) and ants were allowed to forage on a 1M sugar solution. Pheromone deposition on the uncovered part of section X was monitored until 35-40 deposits were reached, after which foraging was stopped and section X, minus segment Y, was transferred to a second T maze for experiment 2**. C)** Experimental design used for experiment 2. Ants were allowed to locate a 1M sugar solution on a randomly chosen branch. While feeding, ants were marked with a paint dot and allowed to return to the nest. Section X was then transferred to the bifurcation so that marked returning ants were faced with a conflict between their route memory and trail pheromone. Decisions were recorded once an ant passed either of the two decision lines. Naive ants with no memory were also tested to determine their response to pheromone alone, and as a control for any side bias.

### Experiment 1: Does the frequency of pheromone deposition change with light level?

Ants were allowed to locate and feed on a drop of 1M sucrose solution, randomly allocated to the end of the left or right branch of the T maze (Fig. 1A). A 5 cm long section of paper, located just before the branches of the T, was marked by lines at either end, and a video camera (Sony; model HDR-XR520) was positioned to record, from the side, all pheromone laying behaviour of ants walking along this designated section (Fig. 1A). This section was chosen because ants were observed to regularly deposit pheromone near the junction and for ease of monitoring. The low lux camera setting was used for the moderate light and darkness treatments. Depending on foraging activity of the colony, the first 12-15 ants that reached the food source and began to feed were marked with a dot of grey acrylic paint (the most discernible colour under IR light). All unmarked ants were removed from the bridge and T maze. The marked ants were allowed to find their way back to the nest, return to the food source, and then once again return towards the nest. Marked ants were removed after passing through the observation section on this final trip. Thus, a maximum of 3 journeys were recorded for each ant: the first return to the nest, the first return to the food, and the second return to the nest. When analysing the videos we assumed that an ant deposited a drop of pheromone each time we saw it clearly curve and dip its gaster to the surface [45]. The experiment was carried out under the three different lighting regimes using six colonies.

### Experiment 2: Effect of illumination on information use choice

To test whether reliance on trail pheromones increases at lower light levels, foraging ants were presented with a conflict between their own route memory and a pheromone trail at a T junction. Following [22], a pheromone trail was created by allowing ants to freely forage on a drop of 1M sucrose situated on the T maze before the bifurcation (Fig. 1B). A piece of paper (X; 10 × 2 cm) was placed directly before the food source with a section of it (4 × 2 cm) covered by an additional piece of paper (segment Y). This ensured that the covered section beneath segment Y remained free from pheromone deposited by ants leaving and returning to the food source. A consistent trail strength was achieved by ending foraging once 35-40 pheromone depositions had been recorded. The maximum time allowed for trail establishment was 20 min; if the minimum number of depositions was not reached in this time, the experiment was terminated.

Memory was then established by placing a 1M sugar solution source on the end of a randomly selected branch of the T maze and allowing the ants to find the food source via the bridge. Feeding ants were marked with a dot of grey acrylic paint and allowed to return to the nest. At high motivation levels such as these, 75-80% of *Lasius niger* foragers take the correct arm of an unmarked T maze at normal levels of illumination after only one visit [22,46]. Before these marked ants left the nest to relocate the food, section X was transferred to the bifurcation of the T maze (Fig. 1B) with the pheromone marked side placed on the branch opposite to where the food source had initially been situated. The covering piece Y was removed so that the bifurcation now had two new arms, only one of which was marked with pheromone. The decisions of the returning marked ants were then recorded. The maximum time allowed for memory development and subsequent decisions by the ants was 30 mins, giving a total maximum experimental time of 50 min, when including trail establishment, which corresponds to the mean trail-lifetime reported for *Lasius niger* [36,47]. Decisions were recorded for ants from nine colonies, tested under the three different lighting regimes.

### Experiment 3: Is memory based solely upon visual cues?

The aim of this experiment was to investigate whether ants could develop a route memory in the absence of visual cues. As in experiment 2 the nest was connected to the T maze by a cardboard bridge and a 1M sucrose solution was placed at the end of a randomly assigned branch. In darkness (0.0007 lux), foraging ants were allowed to locate the food source and were subsequently marked with grey acrylic paint while feeding. Unmarked ants were removed from the maze and marked ants allowed to return to the nest. Fresh paper was placed on the T maze to remove any pheromone present and the binary choices made by returning marked ants at the T junction were recorded and compared to those of naive ants. Ten colonies were used in this experiment.

### Statistical Analysis

Data for the pheromone deposition frequency were found to be zero inflated so we consequently chose to use the MCMCglmm package [48] implemented in R v. 2.14.2 [49] using the zipoisson family function. Uninformative prior distributions were used for fixed effect parameters with a mean of 0 and a large variance of 10^8^. Priors for the variance components were inverse-Wishart distributed with the degree of belief parameter (n) set at ¼ 0.01 and variance (V) limited to 1. Each model was run for 120,000 Markov chain Monte Carlo (MCMC) simulation iterations with a burn-in of 40,000 iterations and a thinning interval of 10 iterations. Autocorrelation between successive iterations was low (<0.05). Maximal models were created and non-significant fixed effects were sequentially removed from the model. Models were compared using the deviance information criterion (DIC). The fixed effects included light treatment (levels of bright, moderate and darkness) and journey (levels of towards nest (1 & 2) & towards food source) while colony and date were used as independent random effects. Mean parameter estimates and 95% credible intervals were constructed and are reported in the results; where estimates do not range over zero, the parameter is deemed to be significant.

Data from experiments 2 and 3 were analysed using generalised linear mixed-effect models (GLMM) with binomial errors in R v.3.4.1 [49]. Models were fitted using the lmer function [50]. Following [51], models were constructed based on *a-priori* expectations. Differences from random choice were tested using exact binomial tests.

## Results

### Experiment 1: Does the frequency of pheromone deposition change with luminance?

Overall, as light level drops from bright to moderate, pheromone deposition significantly increased from 0.45 depositions per passage to 0.76 [parameter estimate = −1.477, 95% CI = (−2.769,-0.122); near darkness vs bright; parameter estimate = −1.206, 95% CI = (−2.267,-0.167), Fig. 2]. However, pheromone deposition did not continue to increase when light levels were further reduced from 10 lux to 0.0007 lux [mean 0.73 depositions per passage, parameter estimate = −0.488, 95% CI = (−1.784, 0.884)].

**Fig 2.**
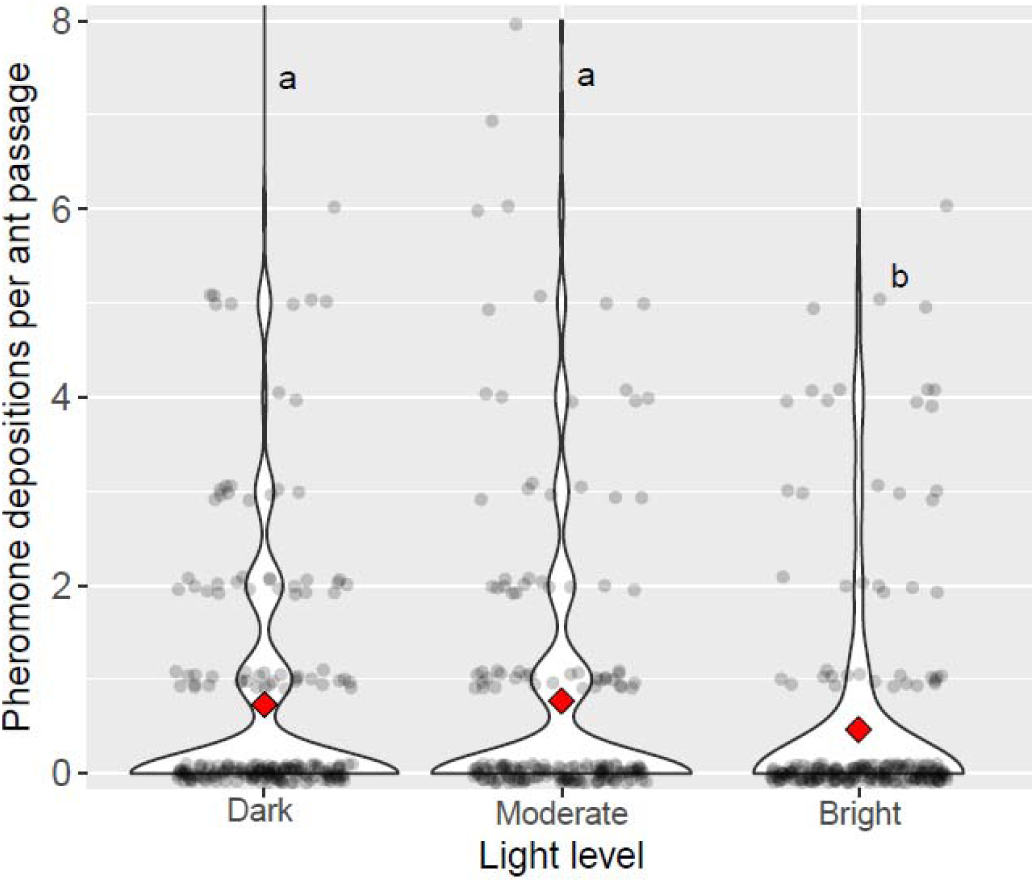
Violin plots show the number of pheromone deposits per passage for the 3 different light levels (dark, moderate and bright) for the three journeys combined. The deposition rate significantly increases when light intensity is reduced to 10 lux, but no further increase is seen with a further reduction in light intensity. Circles are individual data points, red diamonds denote means. Different letters (a, b) signify significant (p <0.05) differences between groups.

While the pattern of increasing pheromone deposition with decreasing light intensity holds over all three visits, the specifics differ. Significantly more deposits were made on the first journey back to the nest under the moderate light level compared to the other two light treatments [Moderate vs Bright; 0.99 vs 0.39, parameter estimate = 2.175, 95% CI = (0.605, 3.713), Moderate vs Darkness; 0.99 vs 0.39, parameter estimate = 2.309, 95% CI = (1.048, 3.602), Fig. 3A]. Of particular note is the significantly greater number of depositions on the return journey from the nest to the food source in the dark, when compared to either moderate or bright light levels [Dark vs Moderate; 0.81 vs 0.41, parameter estimate = −2.249, 95% CI = (−3.509, −0.880), Dark vs Bright; 0.81 vs 0.18, parameter estimate = −2.184, 95% CI = (−3.973, −0.600) Fig. 3B]. On the second return journey to the nest, pheromone deposition was almost one deposition per passage in both darkness and moderate light, but in each case was found not to differ significantly from 0.77 deposits per passage found in bright light conditions [Dark vs Bright; 0.96 vs 0.77, parameter estimate = −0.462, 95% CI = (−2.210, 1.158), Moderate vs Bright; 0.96 vs 0.77, parameter estimate = −0.834, 95% CI = (−2.373, 0.884) Fig. 3C]. There was no significant difference in the rate of pheromone deposition between the two return journeys to the nest in either the bright or moderate light conditions [Bright; 0.39 vs 0.77; parameter estimate = 0.565, 95% CI = (−0.842, 1.730), moderate; 0.99 vs 0.96, parameter estimate = −0.174, 95% CI = (−1.351, 1.019)], but in the dark pheromone deposition significantly increased on the second return journey [darkness; 0.39 vs 0.96, parameter estimate = 1.066, 95% CI = (0.085, 1.929)].

**Fig 3.**
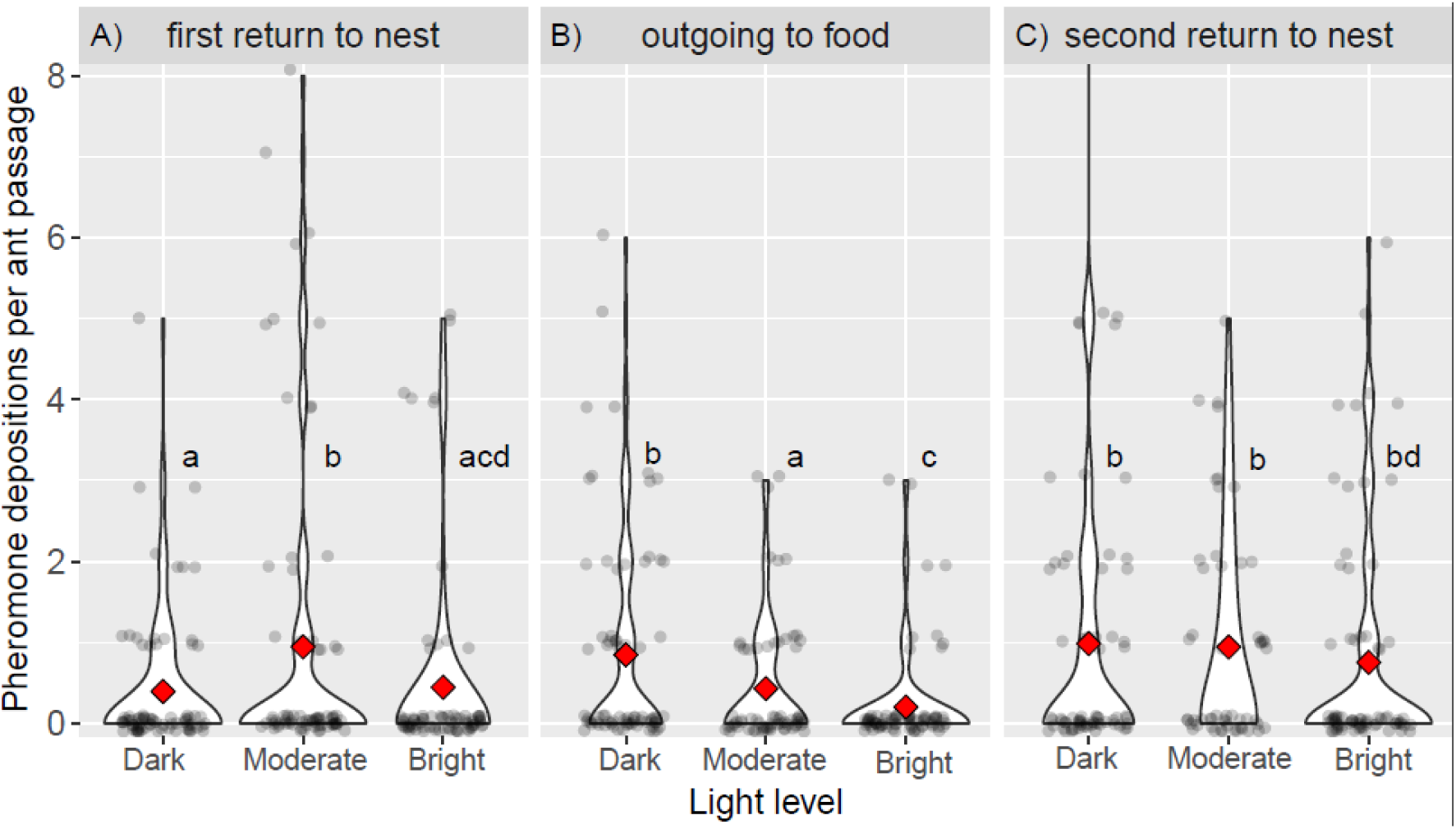
Violin plots show pheromone depositions per passage for the three separate journeys for each of the three light levels. During the first inward journey to the nest **(A)**, deposition frequency is greatest at moderate light levels, while for the subsequent outward journey **(B)**, the deposition frequency is significantly greater in darkness. For the second inward journey **(C)**, there is no significant difference between the deposition rates for three lighting regimes. Circles are individual data points, red diamonds denote means. Different letters (a, b, c, d) signify significant (p <0.05) differences between groups.

### Experiment 2: Effect of illumination on information use

The proportion of ant foragers following the pheromone trail rather than their route memories increases with decreasing light intensity (Fig. 4A). Under bright light only 28 % of ants chose the pheromone treated branch, significantly less than the 61 % seen in darkness (z = −3.56, P = <0.001). More ants followed the pheromone trail in darkness than the 44% in moderate light (z = 1.9, P = 0.059), and in moderate light vs bright light levels (z = 1.78, P = 0.076), but these trends were not significant. Naïve ants followed the pheromone trail at the highest rate, which was significantly more than ants in bright and moderate light levels (vs bright, z = 5.51, P < 0.0001, vs moderate, z = 3.70, P = 0.0002), but not different to ants in darkness (z = 1.45, P = 0.15). The random effect of colony contributes very little to the overall variance (<0.1 %). Ants in the bright light treatment significantly preferred to follow their memories (exact binomial test, 16/59, P = 0.00058). The decisions of ants under moderate light and darkness did not significantly differ from random (moderate light: 25/58, P = 0.36, near darkness: 36/60, P = 0.16). Naïve ants significantly followed the pheromone-marked path (106/148, P < 0.0001).

**Fig 4.**
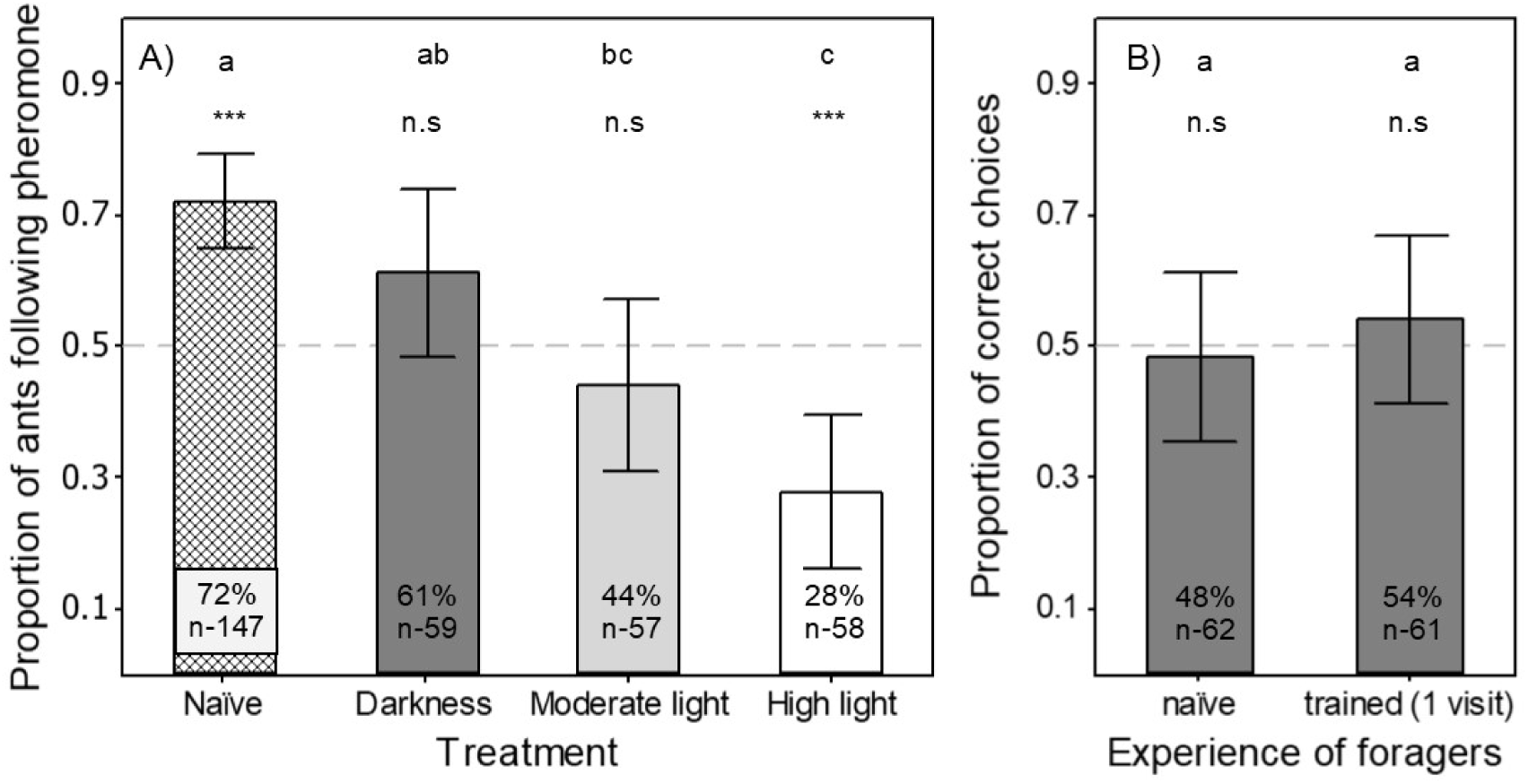
**A)** Histogram showing the proportion of ants choosing the branch of a T maze treated with trail pheromone when presented with a conflict between their own route memory (acquired from one visit) and trail pheromone, under three different lighting regimes (Darkness, Moderate light, and Bright Light). A comparison is also made to naïve ants with no route memory. Response to pheromone significantly increases as light drops from bright to moderate. **B)** Histogram showing the proportion of ants choosing the branch of a T maze that leads to a food source in darkness. Different letters (a, b, c, given above each bar) denote significant differences. *** denotes a group different from 0.5 (P< 0.001) and n.s. denotes not significantly different from 0.5. Error bars represent 95% confidence intervals. Percentages in bars give the percent of ants following the pheromone trail (A) or choosing the arm where food was encountered (B), and numbers denote group sample size. The dashed horizontal lines denotes 50% random choice.

### Experiment 3: Is memory based solely upon visual cues?

Ants which had made one visit to a food source at the end of a T-maze under 0.0007 lux did not perform better than naive ants when choosing a branch at the bifurcation (Fig. 4B). While 48 % of naive ants chose the branch to the food, only 54 % of ants with a memory made the correct decision (z = 0.99, P = 0.31). The random effect of colony and date contributed very little to the overall variance (<0.1 %).

## Discussion

Light levels have a large effect on the way in which *L. niger* foragers make use of, and deposit, pheromone trails. As previously reported [19,22,30], when route memories (private information) and pheromone trails (social information) conflict in bright light (3200 lux), *L. niger* foragers mostly follow their route memories. However, we found that as light levels decreased to dusk-like levels (10 lux) and on to darkness (0.0007 lux), the rate at which ants relied on private information decreased, until in darkness over half the ants chose to follow a pheromone-marked branch. This shift in cue reliance has been previously reported in field observations on two Formica species, *F. polyctena* [31] and *F. nigricans* [52], but our study is the first to demonstrate this under stringently controlled laboratory conditions. The shift from private information use to a reliance on pheromone trails, which we and previous researchers report, is consistent with ants following a ‘copy when uncertain’ strategy, with reliance on private information decreasing as its information content decreases. ‘Copy when uncertain’ is an adaptive information use strategy in many situations, and is employed by vertebrates in a variety of situations [2,53–55]. Recently, behaviour consistent with ‘copy when uncertain’ has been described in the behaviour of *Temnothorax* rock ants during house-hunting, where informed ants rely more on social information about nest quality when their private information is uncertain [33]. Bumblebees in a foraging context have also reported to ‘copy when uncertain’, being more likely to land next to bee models in uncertain environments [32]. Ants have also been reported to shift their reliance on visual and non-social olfactory cues in response to changes in light levels – when trained to locate food using both olfactory and visual cues, various *Myrmica* species preferentially follow visual cues when olfactory and visual cues conflict. However, at lower light levels their preference shifts towards a reliance on olfactory cues, in some cases even when light levels are at an moderate level of 110 lux [56–58].

However, it is noticeable that trained ants follow pheromone-marked paths less than naïve ants in darkness. While not significant, this is puzzling. The rate of pheromone following in naïve ants agrees well with previous data [22,59], but the pheromone following rate of ants trained in darkness is lower, and not different from chance. Why is this? As ants seem unable to learn location after one visit in darkness (Fig. 4B), it seems unlikely that this is due to a conflict between two information sources. An alternative possibility is that they are in a wrong-task state [59,60]; naïve ants may be actively scouting (exploring or otherwise ready to make use of social information), while experienced ants may be in a foraging state, attempting to exploit private information even if it is not there, and disregarding social information. However, Czaczkes et al. [59] showed that task state does not in fact influence pheromone following. It may therefore simply be that the apparent difference between these two groups is not real, and that the 61% of darkness trained ants following pheromone is an underestimate of the true rate of trail following. Similarly, although more than half of the ants in darkness chose the pheromone-marked arm, this was not significantly different from chance rates. We believe this to be a false negative (type II error). If it is not, however, then our results must be interpreted not as evidence for a ‘copy when uncertain’ strategy, but rather as ‘innovate when uncertain’ – ants demonstrably had access to the pheromone information, but may have chosen not to use it.

Ants do not seem to be able to learn a location in the dark after one visit (54% correct choices), while they are very capable of doing so on almost identical mazes in lit conditions (c. 75% correct choices, [22,46]). It is possible that, given more visits, ants would learn to navigate the maze using ideothetic cues, as has been shown in other ant species [61]. It is worth noting that visual cues are in principle not required for navigation by path integration [62], where an odometer linked to any directional cue can be used to estimate displacement from a starting location. Magnetic cues have been shown to be used for navigation by several animals, including ants, especially when other cues are unavailable [63,64]. As path integration is usually used as the initial navigation mechanism, before route-based navigation memories are formed [35], it seems that *L. niger* require visual directional cues to perform path integration.

Pheromone deposition in *L. niger* is very variable between workers, with most ants depositing nothing, and some depositing many dots per passage. However, as pheromone depositions are cumulative, it is the mean pheromone deposition rates that are relevant to the colony. Inspection of these showed that rates of pheromone deposition varied strongly with light levels. Broadly, over all visits, ants deposited about 40% less pheromone at the bright light than in the moderate light level or darkness (Fig. 2). This supports the assertion made above that foragers are less confident of their location in moderate light levels and darkness. Ants which go on to make a navigational error, deposit less pheromone than ants which will make a correct choice [15], further suggesting that pheromone deposition rates are linked to navigational certainty.

A more complex picture emerges when we examine each journey of the ants separately. On the first return to the nest, ants in moderate light deposit about 45% more pheromone than ants in darkness, and also 20% more than ants in bright light (Fig. 3). We interpret this again in terms of certainty; ants which make an error and then correct themselves increase their pheromone laying rate [15,16], presumably in order to assist themselves and their sisters by providing more information for apparently hard-to-learn routes. Thus, ants in moderate light can be interpreted to sense that more information is needed compared to ants in bright light, and therefore provide this information. Ants in darkness deposit very little pheromone, but this is not surprising – ants which are lost or unexpectedly leave a pheromone trail deposit little or no pheromone (65, TJC pers. obs). Surprisingly, on their return to the food source, outgoing ants in darkness deposit on average almost 78% more pheromone than ants returning in bright light. This was unexpected, given that they apparently cannot know exactly where they are going (Fig. 4B). However, as some pheromone and home-range markings had already been deposited, this may act as a reassurance that ants are on the right path [6,65,66]. Given that they are on the right path, reinforcing the pheromone signal provides more information in darkness, where visual information is lacking. Finally, on the second return to the nest, ants consistently deposit a high amount of pheromone at all light levels. We interpret this as all ants, having found food twice in quick succession, being confident enough of their location to recruit strongly to the food source.

How animals strategically use and deploy information has been the subject of intense research [1,2,4]. The information richness of an information source is predicted to be a strong driver of information source use [14]. Studying visual information use under varying light levels provides a powerful and ecologically relevant means of manipulating information richness. By taking advantage of this, we show that information use during ant foraging is consistent with a ‘copy if uncertain’ strategy. We also demonstrate that ants vary their provision of an alternative information source as their primary information source becomes less informative.

## Author contributions

SJ and JB conceived of the study. SJ planned the study and performed the statistical analysis. SJ, AG and EG collected the data. TJC, SJ and JB wrote the manuscript. All authors gave final approval for this work.

## Acknowledgements

Thanks to Stephanie Wendt for help with the violin plots. SJ was funded by the British Biotechnology Research Council (BBSRC). TJC was funded by a Deutsche Forschungsgemeinschaft Emmy Noether grant number CZ 237/1-1.

